# Facial Micro-Movements as a Proxy of Increasingly Erratic Heart Rate Variability While Experiencing Pressure Pain

**DOI:** 10.1101/2025.09.09.674997

**Authors:** Elizabeth B Torres, Mona Elsayed

**Affiliations:** Sensory Motor Integration Laboratory, Psychology Department, Rutgers the State University of New Jersey, New Brunswick, NJ; Rutgers University Center for Cognitive Science, Rutgers the State University of New Jersey, New Brunswick, NJ; Center for Biomedicine Imaging and Modelling, Computer Science Department, Rutgers the State University of New Jersey, New Brunswick, NJ

**Keywords:** pain_1_, facial micro-movements_2_, HRV_3_, stochastic process_4_, proxy parameter_5_

## Abstract

The sensation of pain varies from person to person. These patterns of individual variations are difficult to capture using coarse subjective self-reports. However, they are important when prescribing therapies and tailoring them to the person’s own sensations. Pain can be experienced differently by the same person, and fluctuate differently depending on the context, yet most analyses treat the problem under a one-size-fits-all model. In this work, we introduce a series of assays to objectively assess pressure pain across tasks with different motoric and cognitive demands, in relation to resting state. In a cohort of healthy individuals, we examine pain-free *vs*. pain states at rest, during drawing with heavy cognitive demands, during pointing to a visual target, and during a grooved peg task like inserting a grooved key in a matching grooved keyhole. We recorded Face videos, electrocardiographic signals and adopt a standardized data type called the micro-movement spikes (MMS) to characterize the biorhythmic activities of the Face micro-expressions and of the micro-fluctuations in the heart’s inter beat interval timings. Using the MMS peaks, we find that the continuous Gamma family of probability distribution functions best fit the frequency histograms of both the Face and the heart data. Further, we find that the Gamma shape and scale parameters in both signals span a scaling power law whereby as the noise- to-signal ratio (Gamma scale parameter) increases, so does the randomness of the stochastic process (the Gamma shape decreases towards the memoryless exponential range). We find that as the heart IBI turns more erratic (noisier and more random) the facial ophthalmic region increases the noise and randomness too, with higher linear correlation for tasks requiring haptic feedback (R^2^ 0.84) than for tasks requiring higher cognitive and memory loads (R^2^ 0.77). Increases in transfer entropy shows that recent past activity (∼167ms back) of the heart IBI and Face combined lower the uncertainty in the prediction of the present ophthalmic-Face activity, suggesting that this Face region may serve as a proxy of an increasingly dysregulated heart. These results bear implications for the detection and monitoring of pressure pain and heart dysregulated states.

**Scope Statement:** This work uncovers personalized thresholds of pain through the combination of the participant’s fluctuations in HRV and facial micro-expressions. These signals are obtained before and after experiencing the sensation of pressure pain compared to a pain-free control condition. Tasks with higher haptic demand evoke higher differentiation in the signals than tasks with higher cognitive and memory loads. As the detection and tracking of such individual thresholds is possible through these unobtrusive and highly scalable methods, the work has implications for the personalized tailoring of pain treatments beyond the realms of the lab.

## 1 Introduction

The sensation of physical pain can be experienced through different afferent channels and reach a conscious level, as one attempts to describe it amidst the constant flow of kinesthetic reafference that movements and biorhythms across the body generate. How can the brain’s self-generated biorhythms then differentiate pressure pain, in relation to the body’s resting state and produce a conscious recollection of the experience? We know from the literature that reports of pressure pain are often associated with individuals who experience chronic pain (1), while such individuals seem to have a lower pressure pain threshold than healthy controls (2). Often, these reports are provided by the person while at rest. However, in the context of activities of daily living and natural situations where pain is present, or has been experienced, it may be pertinent to ask if biorhythms from self-generated movements across a variety of tasks could differentiate between the tasks before and during experiencing a level of pain high enough to reach conscious recollection of it.

One channel that may be amenable to discern intensity in physical body experiences and differentiate pain sensations amidst natural motions embedded in naturalistic tasks, could be the Face. Facial micro-expressions can transpire largely beneath awareness and yet, help us automatically differentiate transitions across behavioral states that are now capturable with simple means, like a webcam or a cell phone app using the phone’s camera (3). With the advent of new models from computer vision, it is possible to convert video of the Face to a Face grid and register minute fluctuations in facial micro- motions. Using new algorithms from our lab, one can then extract movement-relevant information within very brief (5 seconds long) videos. Such methods offer the advantage of sampling with ease and do so frequently across a multiplicity of contexts, including the comfort of one’s home. If we measure other biorhythms in tandem with those from the Face, we can also characterize internal physiological activities informing us of possible states of distress and anxiety, such as those present during the flight-or-fight mode of the heart’s biorhythms (4, 5). Measuring the heart rate variability (HRV) across different states of pain, relative to rest, could then be a way to add information to Face data and approach pain differentiation through a multitude of reafferent channels. The autonomic channels are particularly suitable for this as the field of cardio-physiology has made large strides in the analysis of HRV data (e.g., (6–8)). Their results can serve as ground truth to derive, using the Face video-data, highly scalable, unobtrusive means offering proxy signals of pain sensation as a metric of autonomic dysregulation.

In this work, we examine the HRV and the facial micro-expressions before and during the lingering sensation of pain, experienced when performing different tasks with different levels of kinesthetic reafference and cognitive/memory demands. From purely continuous movement speed feedback to movement speed feedback combined with haptic- and pressure-based feedback, we here characterize levels of noise in both the HRV and the facial micro-movements and quantify the degree to which some tasks help us differentiate the experienced pain sensation better than others. We find that shifts in the noise patterns of the heart and the Face are linearly correlated in tasks with haptic-pressure feedback demands. These tasks also result in higher pain differentiation than tasks that have additional cognitive/memory demands, suggesting that cognitive/memory demands could be used as pain distractors. Further, we found in such tasks, evidence that recent past signals of the heart inter-beat interval timings – during more erratic pain activity – tend to increase the certainty of prediction of the present Face micro-motions. We propose that the moment-to-moment micro-fluctuations in facial micro-movements during heart dysregulation may serve as proxy of such erratic autonomic patterns. The implications of this research for scalability and pressure-pain treatments’ design are discussed.

## 2 Materials and Methods

### 2.1 Participants

In this study, 45 healthy, neurotypical adults (27 female and 18 males) of college age were recruited to participate via flyers, advertisements, or through the Rutgers Human Research Pool system. All participants provided informed consent, which was approved by the Rutgers University Institutional Review Board (Study ID #Pro2019000615). Wearable sensors and webcams were used to record biorhythmic activity across the Face, heart, and body while the individual performed each task separately. In a subset of 21 individuals, we recorded concurrent Heart and Face activities and analyzed their performance. Figure 1A-D depicts the tasks in the performed order, while Figure 1E-G provides the assay to elicit the responses across conditions. All tasks were subject to this assay comprising an adaptation phase (denoted pre-pain or control interchangeably throughout the paper and a pain phase where the person experienced physical pressure pain). This work is part of a PhD thesis published in 2024 (5).

**Figure 1.**
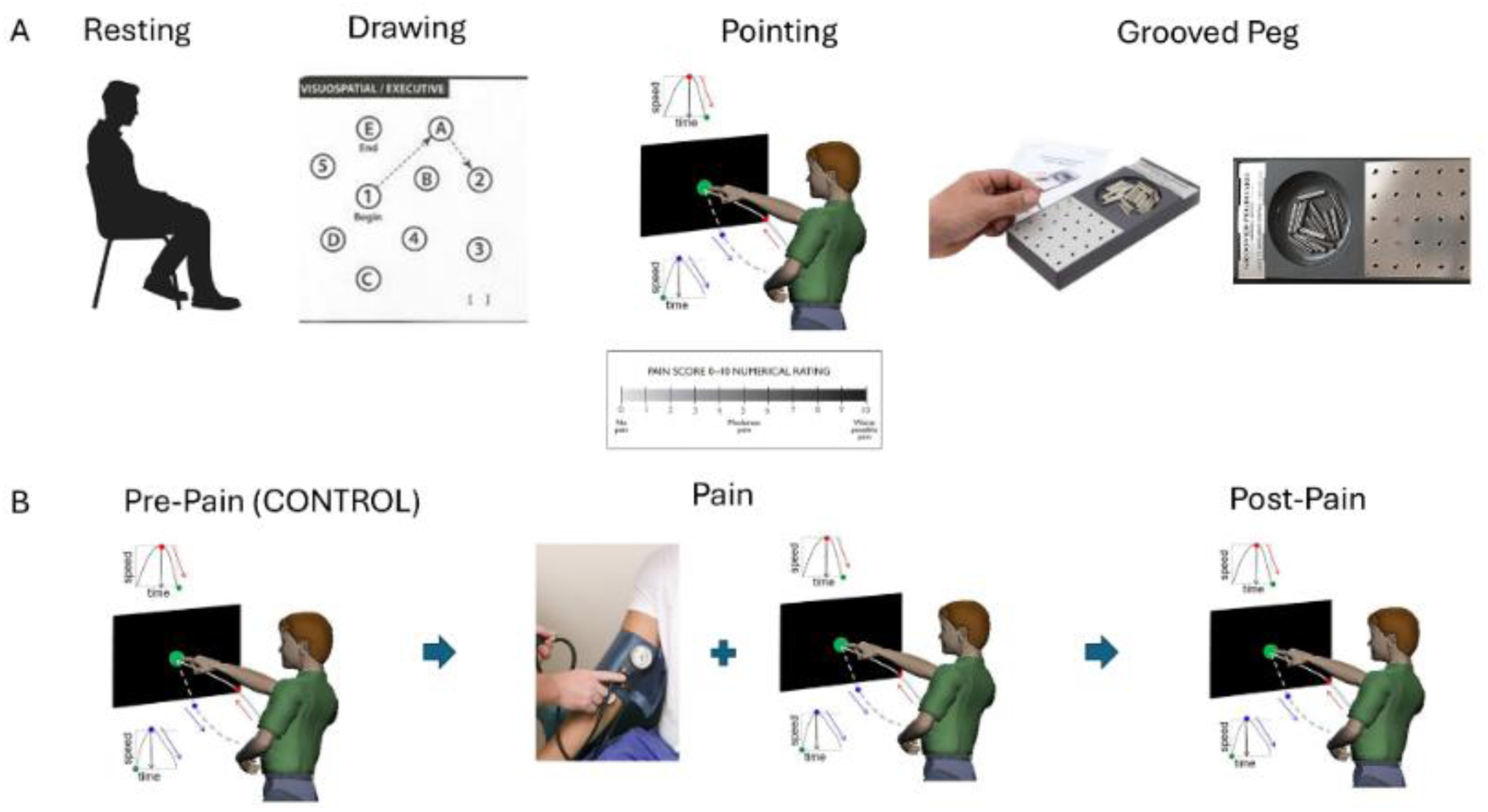
Different tasks to measure the effects of pressure pain. (A) Resting task where the person sits and is instructed to remain as still as possible. Drawing task where the person draws a line connecting digits and letters in a specified order (1 A, 2 B, etc.) from beginning to end. Pointing task, where the person is asked to point to a target (the full loop of forward motion to the target and backward motion, returning the hand towards rest, are captured). The second backward segment occurs spontaneously, without instruction, while the first motion towards the target is instructed. The grooved peg task where the person picks up a grooved peg and aims it at the grooved hole on the board, to insert it along the appropriate orientation. This task requires orienting the grooved peg to match the grooved hole, as when opening a door by inserting a key into the lock. (B) Experimental assay to probe the control condition *vs*. the pain condition. For each task, (using here the pointing task as an example) the person performs the task normally, without any pain (control case). A blood pressure cuff is then placed on the non-dominant arm (the arm that is not performing the task) and inflated to 200mmHg for the entirety of the task to ensure the lingering discomfort producing the sensation of physical pressure pain. The person performs the task under pain (Pain condition) and lastly, the person performs the task after the pain disappears (Post-Pain condition).

### 2.2 Tasks and Assays

The tasks were chosen to evoke movements in different contexts, according to different levels of feedback and cognitive/memory demands acting as possible distractors from the pain. Both the Drawing and Grooved Peg Tasks (called the Peg Task from here onward) had a haptic component mediated by pressure. Yet, the Drawing Task had additional memory and cognitive demands that we discuss below. The pain assay required three blocks, which were used for each task: Block 1 was recorded without pain using two sets of trials. These were aimed at adapting the person to the task. Block 2 was the pain condition, where pain was experienced through sustained pressure applied with a blood-pressure cuff on the non-dominant arm that was not used to perform the goals of the tasks. Block 3 was then registering the biorhythms to assess any aftereffects of pain induction or lingering sensation of pain as it dissipated. In this way, the adaptive nature of the movement and the pain experience could be assessed through continuous adaptive processes.

*Task 1* is the Resting Task. Participants were instructed to sit still and comfortably at rest (Figure 1 Resting). This task of sitting still is rather difficult as the human body, even seemingly at rest, is in constant motion. Breathing, fidgeting and in some cases, involuntary tics are common across the population and often exacerbated by psychotropic medications (9), yet these motions transpire largely beneath awareness. By sampling the bodily micro-movements at rest and while instructing the person to remain still, we are in fact sampling the level of volitional control that the person’s system has over involuntary motions.

*Task 2* is the Drawing Task (Figure 1 Drawing). This is a task commonly used in the Montreal Cognitive Assessment test (MOCA)(10) to evaluate the person’s cognitive and memory abilities as the task goals increase in levels of complexity.. The task consists of drawing a connected path across digits in a random order presented on a piece of paper. For example, from 1 to 10, the order could be 2, 5, 3, 7, 4, 1, 9, 10, 6, 8, spread out across the board, and the person would have to join the numbers in order – 1 followed by 2, followed by 3, *etc*. all the way to 10, by continuously tracing the path of the pen on the tablet/board/piece of paper. This action provides continuous feedback from the surFace of the table, based on the pressure that the hand exerts on the pen and the pen on the table ’s surFace. Such continuous feedback provides a type of haptic information that the brain needs to continuously monitor and modulate the applied pressure (needed to keep the pen moving against the surFace). This occurs simultaneously with the cognitive load that the ordering of the digits requires on the space of the paper. The task also taps onto the spatial memory capacity to hold the proper order already visited and the upcoming digit at a different location of the paper. An additional layer of cognitive load emerges when the person must sort an alphanumeric string as the letters A,B,C,D, …, *etc*. are added to the numbers. In our present scenario, 1A, 2B, 3C, …, *etc*., must be considered in the proper ordering to trace the paths connecting the digits and letters randomly located across the paper. Often, the timing of these traces increases when the alphanumeric case is included in the assay (11, 12) evidencing (among other kinematic parameters) that the cognitive and memory loads are much higher in alphanumeric mode than in the simpler digits mode. This task is an example of the intersection between cognition and motor control (12).

*Task 3* consisted of pointing to a visual target (Figure 1 Pointing). This Pointing Task involves a forward, goal-directed phase whereby the hand is deliberately aimed at pointing to the visual target at one’s own pace (comfortable speed). This motion is followed by an uninstructed, spontaneous backward retraction of the hand, bringing the hand back to rest, without even realizing that the motion is taking place. The backward motion is quite automatic. Two fundamental types of movements are coordinated and sequenced throughout the task (13). Previous work with deafferented case study Ian Waterman(14), and autistic participants (15) support the choice of task to probe the sense of kinesthetic reafference and the role of noise signatures on motor control and coordination. Here we were interested in the differentiation of this task in Pain-free *vs*. during Pain condition.

*Task 4* consisted of inserting grooved pegs on a grooved pegboard (Figure 1 Grooved Peg). This Peg Task has a transport phase from the hand resting position to the grooved pegs reservoir, then a phase of picking up the grooved peg (sensing through haptic feedback the groove of the peg and applying adequate pressure to hold it up against gravity) then lifting it to transport it to one of the grooved holes on the peg board and successfully inserting it on the groove-matching hole. This task has a strong biomechanical compliance component and a strong cognitive component as well. The biomechanical component must align the arm and hand according to physical space constraints, to facilitate the execution of the task. The cognitive component must perform multiple coordinate transformations from visual to proprioceptive spaces before the motion initiates and must continuously adjust them during the action execution. A solution to Bernstein’s degrees-of-freedom problem is needed here (16), to translate the groove orientation correctly into the proper configuration of the arm rotational joints that will result in the optimal hand orientation (17). Facilitating the alignment of the grooved peg that the hand sustains with the grooved hole on the pegboard is like successfully inserting a key in a lock, *e.g.*, to open a door. Continuous haptic feedback can then help the brain figure out the proper coordinate transformations necessary to transport and accurately fit the peg in the hole, according to the groove. Modulating the linear speed during translation of the arm limb segments, including the hand, and monitoring the angular speed during rotations of the arm’s joints, occur simultaneously in such goal-directed task during its orientation-matching phase (18). Part of our hypothesis is that, because of their complexity, these mental operations may have the potential to distract the mind from the underlying pain sensation, as the person attempts to solve the task correctly. Multiple attempts / errors are allowed since the goal of our study was to evaluate the nervous system pre- and during-pain, rather than evaluate the accuracy of the performance. Together, the Pointing, Grooved Peg, and Drawing tasks constitute our assay to digitally measure pre- and during-pain sensations, relative to the baseline resting state.

### 2.3 Sensors and Data Extraction

We used videos and wearable sensors to record the facial and bodily biorhythmic data respectively. Below we describe the data acquisition process to motivate the data analysis.

#### 2.3.1 Face Signal Registration

A webcam (as the one depicted in Figure 2A, Logitech, 30Hz) was used to acquire brief videos across the 4 tasks described above and for the pre- and during-pain conditions. The Face projected its natural expressions as the participant engaged in the task. That is, we did not ask the participant to make any facial expression but rather recorded the Face as the tasks and assays unfolded. Upon collecting video, we ran Open Face (see Notes 1 https://github.com/TadasBaltrusaitis/OpenFace/wiki) and retained the facial grid, 3D gazes, and action units. These are shown in Figure 2B. We then used the trigeminal nerve (cranial nerve V) regions, to parse out three subregions of the Face (ophthalmic, maxillary, and mandibular), which we coin V1 colored red, V2 colored blue and V3 colored black, respectively. These are shown in Figure 2C. The sub-regions are important for sensing movement reafference throughout key facial points involved in social and emotional communication (*e.g.*, the eyes, the ears, mouth, lips, tongue, mandibular regions, *etc.*) and offer the potential to differentiate activities according to levels of pain and task complexity.

**Figure 2.**
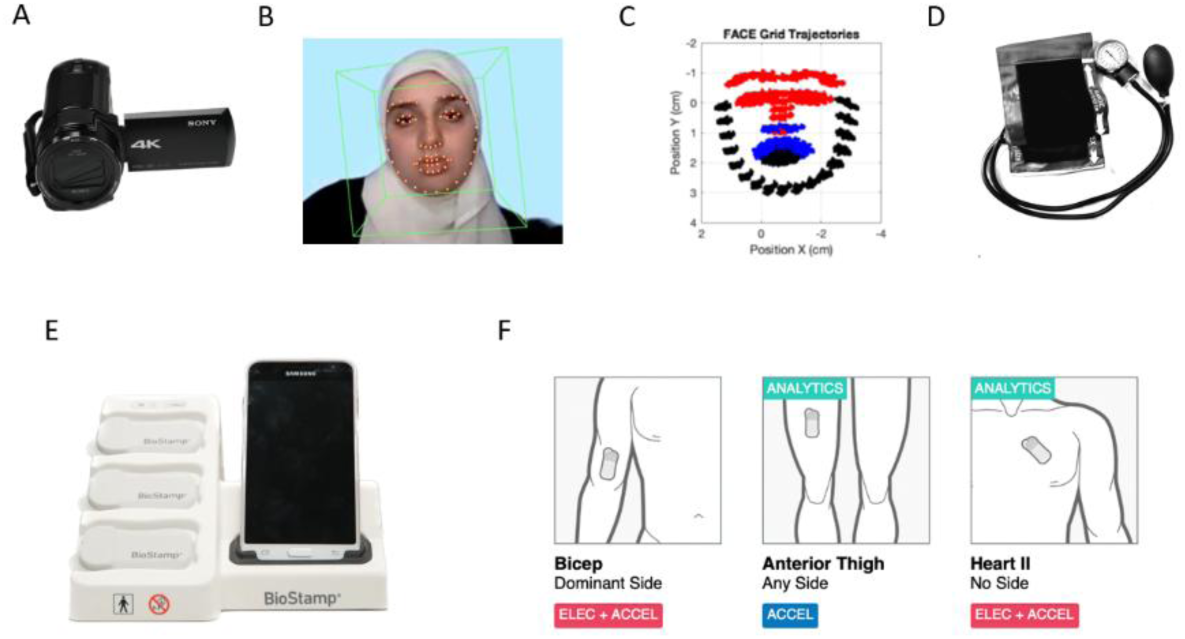
Data acquisition tools. (A) A webcam collects video of the person for the full duration of the task. (B) Open Face software is used to estimate the facial grid from the video of one of the authors (Dr. Elsayed), the 3D gaze and the head orientation, saving action units as well responsible for the micro-motions of the facial muscles. (C) Trigeminal subdivision based on the facial nerves innervating three main regions: V1 (ophthalmic), V2 (maxillary) and V3 (mandibular.) The facial grid is subdivided into V1 (red), V2 (blue), and V3 (black) subregions to study the stochasticity of the micro-motions. (E) Blood pressure cuff used to evoke the sensation of pressure pain in the non-performing arm. (F) The MC10 Biostamp sensors and phone used to collect time series data by registering different biorhythms. (G) Locations of the body where the Biostamp sensors were located to co-register data form the bicep, anterior thigh and heart (image acquired via registered study on MC100 Biostamp-n-Point Cloud system, now acquired by Medidata).

#### 2.3.2 Heart Signal Registration and Pain Induction Procedure

We measured the activity of the heart using the MC10 Biostamp-nPoint system (Lexington, MA), placing one sensor with 4 electrodes on the lead II position of the chest. The blood pressure (standard tourniquet) cuff in Figure 2D was used to measure the amount of pressure sustained (200 mmHg for the entirety of each task ∼ about 1-3 minutes each) to induce the lingering sensation of pain. This method serves as a modified version of the submaximal effort tourniquet test (19) found to emulate pathological pain (20). The Biostamp sensors are depicted in Figure 2E along with the smart phone that enables time stamping the signals, labeling, and automatically uploading them to the cloud. Figure 2F shows the arm location where the Biostamp sensor was affixed. Adhesive tapes with gel were used to help register the electrical activity from the muscles (electromyography EMG) and the heart (electrocardiography ECG). In this paper we focus on the ECG signals and reserve the other data for a different publication.

### 2.4 FACE Data Analysis

We focus on video data acquired from the Face and the inter-beat-intervals’ times acquired from the ECG providing the R-R peaks. Figure 3 provides the analytical pipeline for the Face data while Figure 4 provides it for the heart data. We reserve the eye gaze and head orientation data from the Face videos for a different publication.

**Figure 3.**
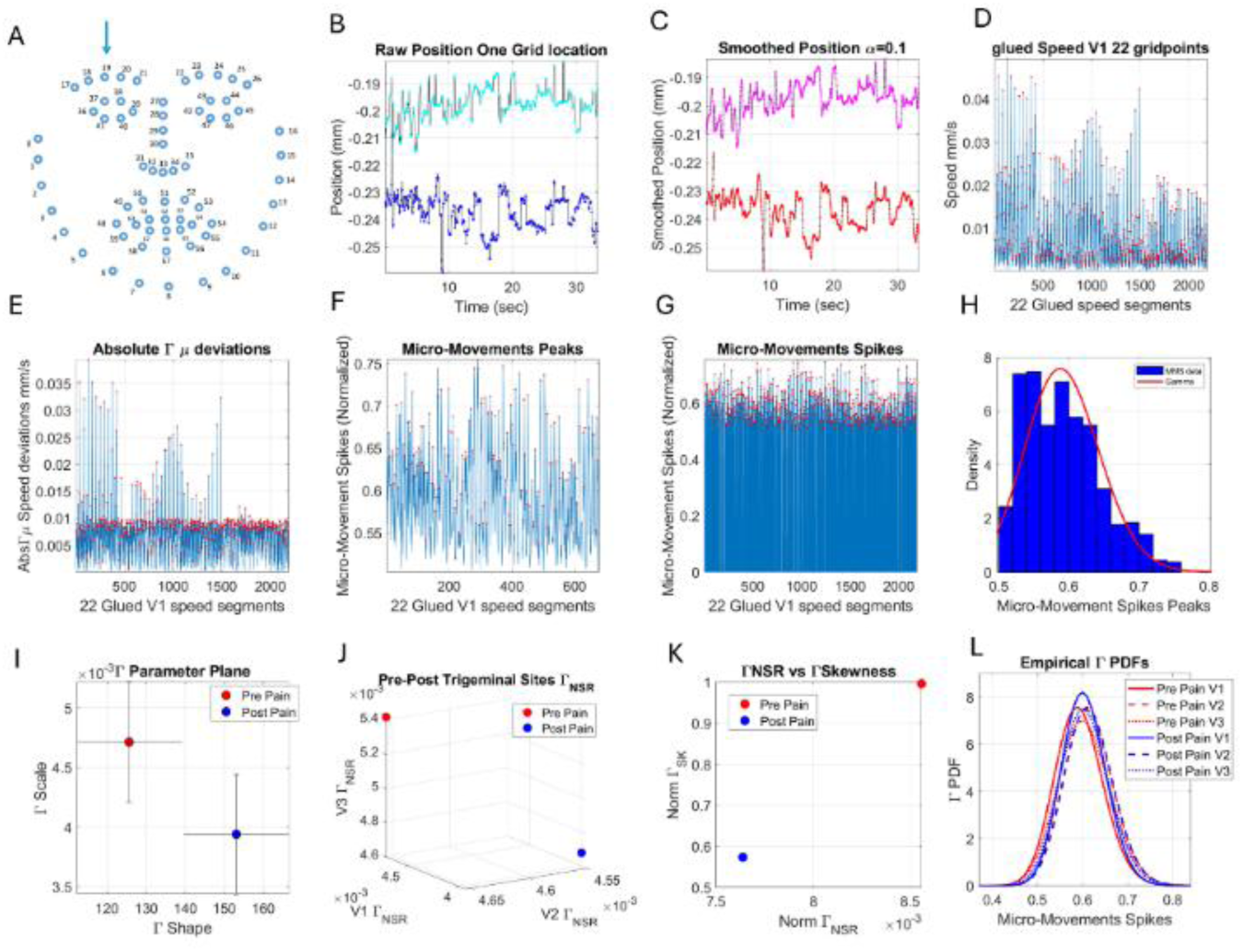
Analytical pipeline of facial biorhythms. (A) Face grid mapped (indexed) across regions ophthalmic V1, maxillary V2, mandibular V3. (B) Sample XY-position traces of one grid point (19 in A) registered for 35 seconds. (C) Smoothed trajectories. (D) Scalar speed profiles from 22 sites in region V1. (E) Absolute speed deviations from empirically estimated mean (from the red peaks). (F) Locally scaling allometric effects due to anatomical disparities in lengths, impacting speed ranges across participants. (G) Full micro-movement spikes conserving the frame of each original speed peak. (H) Frequency histogram of the MMS peaks. (I) Gamma parameter plane used to represent the empirically estimated Gamma PDFs as the shape and scale parameters with 95% confidence intervals for each parameter. (J) Parameter space spanned by the Gamma NSR (scale parameter) of each PDF thus estimated for each Face region. (K) Parameter plane is defined by the norms of the Gamma NSR vector spanned by the V1, V2, and V3 NSRs and the corresponding Gamma PDF skewness. (L) Empirical Gamma PDFs for each region and condition (control *vs.* pain.)

**Figure 4.**
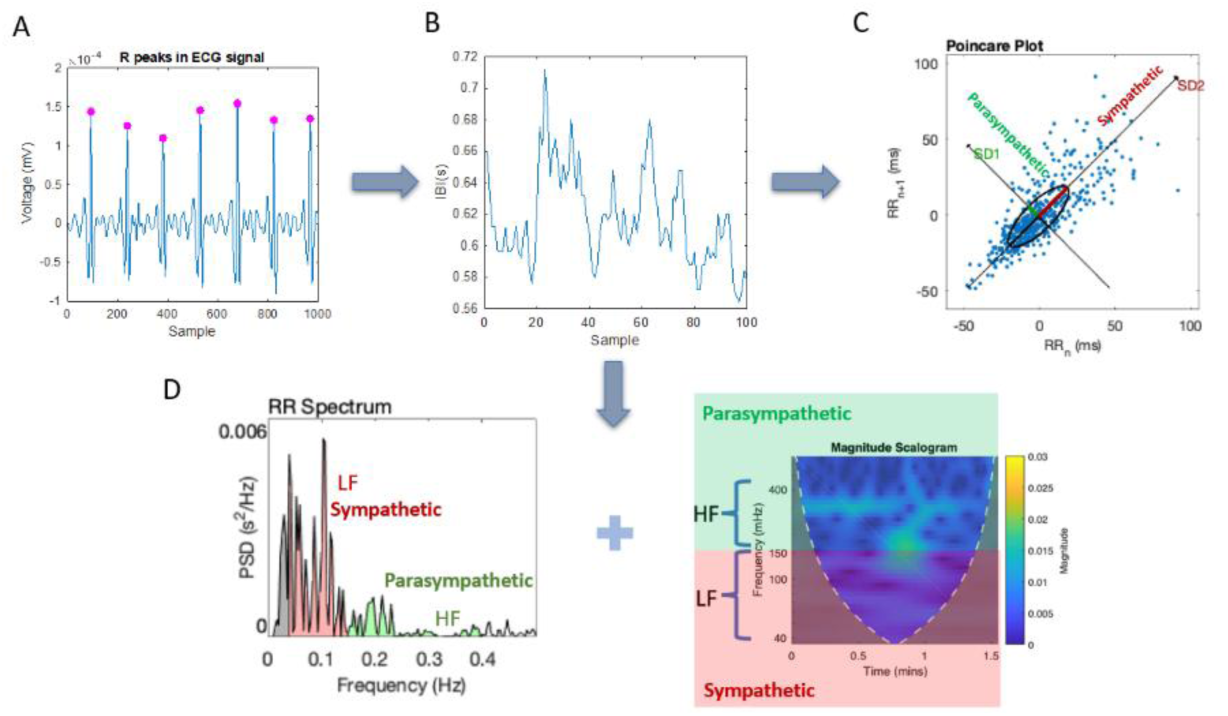
An analytical pipeline for pre-processing the cardiac data. (A) Upon filtering noise and detrending the ECG data, the R-peaks are localized and in (B), the temporal distances between the peaks are obtained. This produces the RR-peaks series data or inter-beat interval times series (IBI). (C) The temporal domain is assessed by obtaining the Poincare plot, a shifted version of the time series one step forward along the y-axis and the current time version along the x-axis. The scatter is fit by an ellipse and the principle axes examined as SD1 (variability associated with the parasympathetic system) and SD2 (variability associated with the sympathetic system). These parameters are then examined in relation to noise regimes from the empirically estimated micro-movements spikes (micro-peaks) distributions. (D) In this step, we perform the frequency domain analysis using power spectral decomposition techniques and obtained the low and high frequency bands for examination of the sympathetic and parasympathetic regimes associated with the lower and higher frequencies, respectively. A magnitude scalogram can then be used to visualize the data as frequency over time with power information (color bar representing the magnitude/power levels).

The Face grid consists of 68 points indexed across the Face in Figure 3A, while considering trigeminal areas V1, V2, and V3 depicted in Figure 2 C. The positional pixel trajectories of facial movements were obtained using a webcam and sampling a few seconds (*e.g.*, 35 seconds are shown in Figure 3B from the location indexed as #19 in Figure 3A). This positional trajectory along the *x* and *y* coordinates was smoothed in Figure 3C using in-house developed code employing spline interpolation routines from MATLAB (version R2023b). The speed profiles derived from the velocity field at each indexed point of each region of the Face were pooled to empirically estimate the probability distribution best fitting the histogram of peaks and then obtaining the first moment (the mean) to subtract the deviations from it. The speed profiles pooled across 22 grid points in V1 are shown in Figure 3D. The positive (absolute) speed deviations from the empirically estimated Gamma mean are shown in Figure 3E. These positive deviations from the mean were normalized by Equation 1 to locally scale out allometric effects due to anatomical disparities in length impacting the speed, our parameter of interest. Accordingly, for higher speed values, on average, the moment-to-moment fluctuations in facial speed will be lower than for slower speed values.

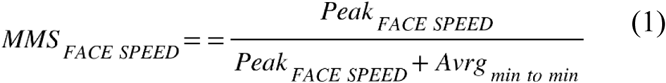

The continuous Gamma family of probability distributions fitted the histogram of speed peaks best, in a maximum likelihood estimation (MLE) sense. The two Gamma parameters, the shape and the scale (dispersion) were estimated from empirical data with 95% confidence. The peaks of the absolute deviations were normalized to eliminate the allometric effects due to disparate anatomical lengths of the Faces across participants. These scaled peaks are shown in Figure 3F for peaks deviating from the empirical Gamma mean. Figure 3G shows the micro-movement spikes in their entirety, preserving the frames where they occurred. Notice that 0-values represent the mean motion and non-zero values are the deviations from the mean. Figure 3H provides an example of a histogram from the micro-movement spikes (peaks) and the best fitting probability distribution function (which also resulted in the continuous Gamma family, according to MLE). Figure 3I shows the Gamma parameter plane with two distribution samples from the control condition (pre-pain) and the pain condition. These points represent the shape and scale parameters of the Gamma PDF estimated for each case, representing a region of the Face. Points are represented by 95% confidence intervals. Figure 3J represents the points for each Face region along the V1 (*x-axis*), V2 (*y-axis*), V3 (*z-axis*) separating the control from pain distribution values of the Gamma PDF’s noise to signal ratio (NSR).

The Gamma NSR is the scale parameter denoting the dispersion of the distribution. Given the Gamma shape *a*, and the Gamma scale *b*, we can see the NSR or variance-to-mean ratio, Gamma variance divided by the Gamma mean, gives the scale parameter *b* as our predictor of the NSR. We have found a strong linear correlation between the log-log of these two parameters spanning several decades in the logarithmic 10-based scale across the human lifespan (15). This result has extended to multiple biorhythms of the nervous systems, thus enabling us to reduce the two parameters to one of interest, since given one parameter, we can infer the other with high certainty. Accordingly, we here examine this scaling power law for the Face and for the Heart data. Further, we examine the Gamma NSR for each Face region and plot it as in Figure 3J, spanning a vector representation where the points are measured from the (0, 0, 0) origin using the Euclidean norm. Based on this representation of the Gamma NSR across the facial regions, we then obtain the scalar value summarizing the facial noise, which also gives us a sense of the randomness of the process under investigation (by inferring the values of the Gamma shape parameter). At one extreme, at a Gamma shape value of 1, the process is memoryless exponential, whereby future events are not predictable by present events, and present events are not predictable by past events. On the other extreme, we have Gaussian-like processes, whereby present and past events can be used to predict future events. The continuous Gamma family spans all these different states and serves (in an MLE sense), in the human empirical biorhythmic data, as an accurate representation of the time series (random process) under consideration.

We then pair the Gamma NSR with the Gamma skewness, giving us a picture of the tail of the distribution. This builds a parameter plane to represent the differentiation between the control case and the pain case for each of the four tasks under consideration. Figure 3K shows the parameter plane for the scalar Gamma NSR values obtained using the Euclidean norm of the Gamma NSR vector (as in Figure 3I) corresponding to the ophthalmic, maxillary, and mandibular regions of the Face. The PDFs corresponding to each region and control *vs*. pain condition for this example are shown in Figure 3L.

### 2.5 Heart Data MMS and Gamma Process Analysis

The electrocardiographic (ECG) time series data registered by the MC10 Biostamp sensor shown in Figure 2F and located on the chest area as in Figure 2H (rightmost panel) was used to isolate the R-peaks and obtain the inter-beat-interval times (IBI) series. These are shown in Step 1 and Step 2 of Figure 4. Upon IBI acquisition, the series was subject to frequency and time-domain analysis, also depicted as Step 3 and Step 4 of Figure 4, respectively. The power spectrum decomposition revealed the power of the signal for the different frequency bands over time, thus dividing the frequency ranges into the low and the high frequency band regimes. These have been previously identified in the literature with the sympathetic (low frequency band spanning to 0.04-0.15 Hz) and parasympathetic (high frequency band spanning 0.15-0.40 Hz.) (21, 22). The former is associated with *flight-or-fight* states and the latter with rest-digest states (22).

A visualization method useful to show power across the frequency bands over time, is the magnitude scalogram, shown in Figure 4 with the corresponding sympathetic *vs.* parasympathetic nervous systems subdivisions. Lastly, the time domain analysis involves the Poincare plot, which examines the dispersion of the scatter formed by the RR times (in ms at time t) *vs*. the RR times advanced one frame (at time *t + 1*) fitted by an ellipse that identifies the principal axes of the variance (standard deviations) as SD1 (shorter axis associated with the parasympathetic system) and SD2 (longer axis associated with the sympathetic system) (23, 24). Motivated by prior results correlating the Gamma NSR with the SD2 values (5), we here ask if the change in SD2 and the changes in Gamma NSR of the Heart IBI signal correspond in any way to the changes in SD2 and Gamma NSR of the Face signals. In both cases, we use the MMS described in the previous section adapted here to the IBI.

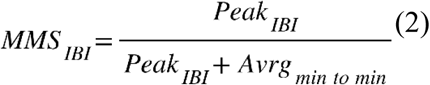

### 2.6 Statistical Comparisons

To evaluate the statistical significance across Gamma distribution parameters, pairwise between conditions, we employed the non-parametric Wilcoxon rank-sum test. In the case of more than two groups, we used the MATLAB routines to implement the Kruskal-Wallis test, a nonparametric one-way ANOVA, extension of the Wilcoxon rank sum test. This test compares the medians of the groups of data in the parameter of interest, to determine if the samples come from the same population. To compute the test statistics, the test uses ranks of the data by sorting it from smallest to largest, instead of numeric values and taking the numeric index of this order. If observations are tied, the rank is equal to the average rank of all observations tied with it. The F-statistic used in classical one-way ANOVA is replaced by a chi-square statistic, and the p-value measures the significance of the chi-square statistic. The Kruskal-Wallis test assumes that all samples come from populations having the same continuous distribution, apart from possibly different locations due to group effects, and that all observations are mutually independent. Both assumptions are held in our data. By contrast, classical one-way ANOVA replaces the first assumption with the stronger assumption that the populations have normal distributions, which is violated in our empirical data.

We followed with the MATLAB function multcompare performing pairwise comparisons through an interactive graph that shows the significantly different means with disjoint (non-overlapping) intervals (25).

### 2.7 Correlations Between Heart and Face Gamma NSR

To try and understand the relationship between the facial biorhythms and the heart’s IBI timings, we used the differences in Gamma scale (the Gamma NSR) from Pain relative to the control (no pain) condition of each task. Linear polynomial fitting R-square measurements and goodness of fit parameters are reported for each task across 21 participants for which both heart and Face data sets were available and taken simultaneously. We used the curve fitting toolbox of MATLAB with bicubic interpolation.

### 2.8 Transfer Entropy Measurements

We treat the two standardized Micro-Movements Spikes (MMS) time series signals derived from the Face and the Heart’s IBI as random processes well characterized empirically by the continuous Gamma family of probability distributions. We empirically estimated these families as they changed for each participant, from task to task, in the control condition. We also empirically estimated these families as they shifted from the control to the pain condition in each task and across all 4 tasks. Then we aimed at further understanding the relationship between these two streams using the notion of Transfer Entropy (TE).

Given process *X* (representing the Face MMS) and process *Y* (representing the Heart’s IBI MMS), the TE is the amount of uncertainty reduced in future values of *Y* by knowing the past values of *X* given past values of *Y*. If *X_t_* and *Y_t_* for, are two random processes and the amount of information is measured as Shannon’s entropy

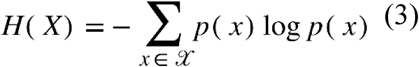

where the denotes the sum over the possible values of variable *x* and we adopt base 2 (unit of bits) to measure information of the variable, then the conditional mutual information with the history of the influenced variable in the condition is denoted:

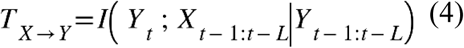

And the TE is denoted:

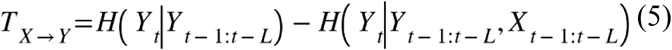

We used the Information Dynamics Toolkit (JIDT) (https://github.com/jlizier/jidt/) infodynamics-dist-1.6 toolbox in MATLAB with open-source code Java implementation, by Joseph T. Lizier, Ipek Özdemir, Pedro Mediano, Emanuele Crosato, Sooraj Sekhar, Oscar Huaigu Xu and David Shorten. We used the continuous-valued TE functions in both directions, by setting *X* in one direction as the Face MMS and *Y* as the Heart IBI MMS, and in the opposite direction, by setting *X* as the Heart IBI MMS and then *Y* as the Face MMS.

We asked if knowing the Face MMS and the Heart IBI MMS in the past could lower the uncertainty of the present Face MMS. We also inverted the question and asked if the current Heart IBI MMS uncertainty decreased given the past Face and Heart IBI MMS. To that end, we used windows of 50 frames with 10% time into the past (5 frames or ∼166.6ms at 30Hz video resolution) for 300 frames representing 10-second-long data from the Face or the Heart IBI states. We note here that despite disparate sampling resolutions between the video capturing and the Biostamp MC10 sensor, the standardized MMS data type brings the two derived signals on a comparable scale of peaks relative to the empirically estimated mean. We can therefore investigate this TE question.

For each task, we sorted the TE values across the 21 participants for the TE obtained in one direction *T_X_*_→*Y*_ and plotted the TE in the other direction *T_Y_*_→*X*_, for each participant. This helped us appreciate the deviations of one curve from the other in those participants where the TE was low *vs.* the cases where the TE was high, relative to the curve of individually sorted values.

*T_IBI Heart_*_→*Face*_ was used as the sorted participants in one comparison, to examine *T_Face_*_→*IBI Heart*_ in both the control condition and the pain condition.

Then, the sorted participants *T_Face_*_→*IBI Heart*_ were used to plot the *T_IBI Heart_*_→*Face*_ values in both conditions.

Lastly, we took the differences in each case, relative to the sorted data, and for each task examined the differences in TE in both directions. We note here that our goal was more modest than establishing causality. We merely ask if knowledge of the recent past in both signals lowers the uncertainty of one of the signals (the Face or the heart, by comparing to one at a time). Since we are measuring this on the shifts in Gamma NSR and we have established the empirical relationship between the Gamma NSR and the distribution shape, we know that decreasing the NSR parallels increasing the shape values away from the memoryless Exponential ranges of the Gamma family. We also know that in other motor biorhythms, this is described by a scaling power law with inverse relation between the Gamma NSR and the Gamma shape – as the NSR decreases (higher signal to noise ratio), the shape shifts to more symmetric regimes featuring more predictive signal and therefore lowering the uncertainty of the system (15). The use of TE here on the standardized MMS streams of the Face and the Heart is then justified by the Gamma NSR serving as proxy of the full distribution, a relation that we have uncovered empirically.

## 3 Results

### 3.1 Tight Linear Relation (Scaling Power Law) Describes the Log Gamma Shape and Log Scale Parameter’s Fit of the IBI MMS Peaks

The frequency histograms of the MMS time series derived from the IBI were well fit by the continuous Gamma family of probability distributions (PDF) which generate for each participant a unique family with personalized shifting patterns. At the cohort level, as with other biorhythms, the scatter of points representing the (Г_SHAPE_, Г_SCALE_) values, whereby each point corresponds to the signature of a participant, also followed a tight linear relationship on the log-log Gamma parameter plane. In this sense, as in other biorhythms well characterized by the MMS Gamma process, knowing one parameter enables us to infer the other. This can be appreciated in Figure 5A-D for each of the tasks and for both the control (red dots) and the pain (blue dots) conditions. The insets of the figure reflect the empirical families of Gamma PDFs derived from the empirically estimated Gamma shape and scale parameters, with 95% confidence. Supplementary Table 1 shows the fitting values, including the slope and intercept values as well and the centered data mean and standard deviation values for each task (Resting, Drawing, Pointing and Peg) and condition (Control *vs*. Pain).

**Figure 5.**
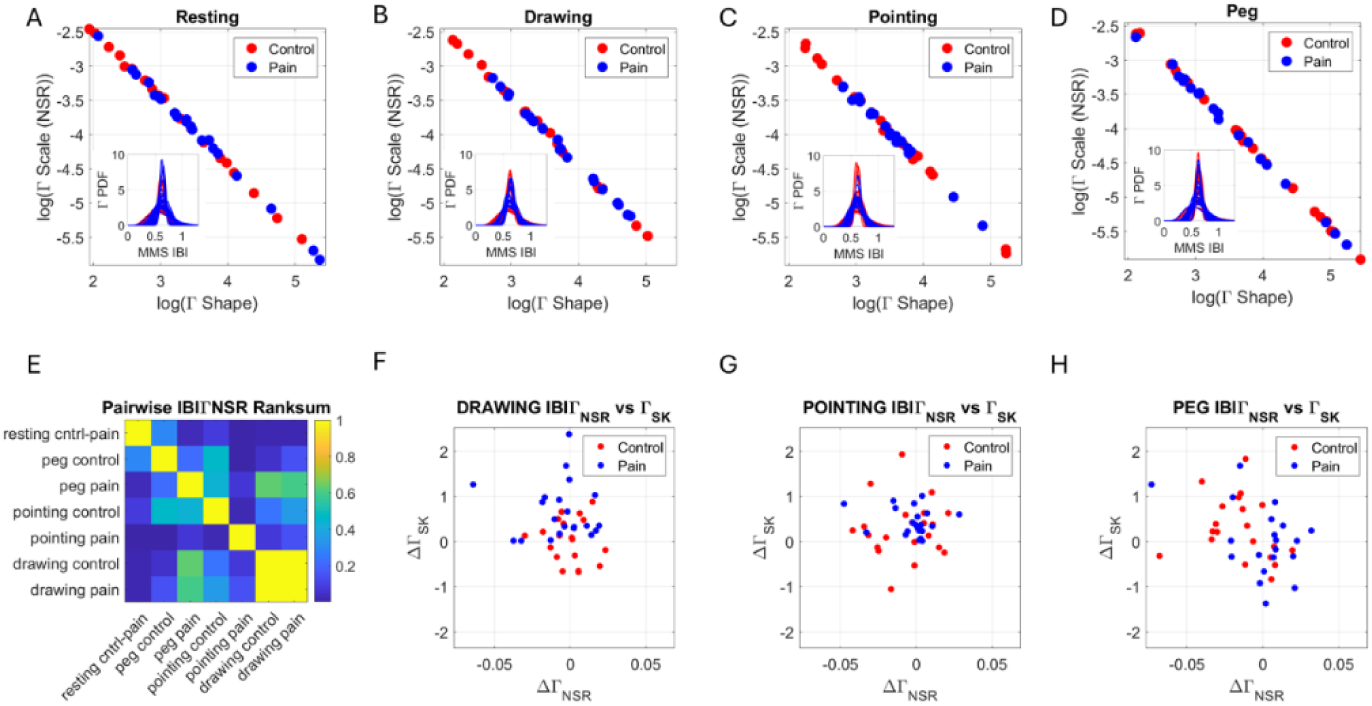
The continuous Gamma family of probability distributions fits well the empirical IBI data in a maximum likelihood estimation sense, with 95% confidence. (A) Control and Pain conditions at resting, (B) Drawing, (C) Pointing and (D) Peg tasks. Each (Г_shape_, Г_scale_) point on the Gamma parameter plane is plotted such that a self-emerging pattern appears, the log-log of the scatter aligns according to a tight linear negative correlation: as the shape of the distribution turns more symmetric (increasing the shape values) the scale (dispersion) values decrease. The scale is the noise to signal ratio (NSR, variance over the mean). (E) The pairwise comparison of the NSR according to the rank-sum test reveals statistically significant differences (*p < 0.01*) for all the tasks during the pain state, relative to the absolute difference between pain and control states for the resting task. (F-H) Parameter plane representing the difference in NSR vs. the difference in skewness for each task relative to the resting task, in both the control (red) and the pain (blue) blocks.

As the shape parameter increases away from the limiting value of 1 (the memoryless exponential most random regime) towards the higher values of heavy tailed distributions and even higher values in the cases of symmetric Gaussian distribution, we see that the scale (dispersion) values decrease. The scale is the dispersion (spread) being also the NSR (as explained in the methods). The negative correlation provided by the tight linear fits to the log-log of the values implies that as the noise decreases, the process becomes more predictable and as the noise increases, the process becomes more random (towards the memoryless exponential regime). The pairwise comparisons of the NSR parameters across participants revealed significant differences (Wilcoxon rank-sum test *p < 0.01*) for the comparison of resting in control *vs*. resting in pain. This was also the case for the other tasks when examining this relative difference (except for the peg during the control block). This can be appreciated along the first row and first column of Figure 5E depicting the color-coded matrix of probability values from the rank- sum test. As the participants performed different tasks, their individualized families of distributions changed shape and scale parameter values. This demonstrates the non-stationary nature of the Heart IBI timings. The shifts from control to pain also demonstrate the susceptibility of this parameter to pain and suggests a good proxy of states of pain.

### 3.2 Heart IBI Gamma PDF Skewness and NSR Shift Across Tasks Relative to Resting State

Given that knowing the Gamma NSR, we can accurately infer the Gamma shape parameter, we turned our attention to this Gamma scale parameter as a proxy of the Gamma PDF spanned by each personalized family (for each participant) and examined it in relation to the corresponding Gamma skewness. To that end, we build a parameter plane whereby points are representing the (Г_NSR_, Г_SKEWNESS_) values. Whereas the scale indicates the NSR (Gamma variance divided by Gamma mean) and provides information on the predictability of the Heart IBI timing, the skewness provides information on the tail of the distribution. At 0 skewness, the distributions are symmetric. Positive values indicate accumulation of larger MMS (standardized deviations from the Gamma mean for the Heart IBI) suggesting shorter interval times between R-peaks, indicating on average faster heart rate and respiration rhythms. Negative values indicate accumulation of smaller MMS suggesting longer interval times between R-peaks, indicating on average slower heart rates and respiration rhythms. This is so because of the standardization of the Heart IBI series by Equation (1). In Equation (1) the term in the denominator indicates the local averages of the signal comprised by the points from local minimum to local minimum surrounding the local peak (maximum). If this value is on average small, the numerator is larger. Thus, shorter Heart IBI timings on average (faster heart rate and respiration rhythms) will produce larger MMS deviations from moment to moment.

For each task, we obtained the differences in NSR and skewness relative to the resting task and appreciated differences in shifting patterns for each task. During the control block, the Drawing Task had a decrease in the skewness values that contrasted with the Peg Task whereby most participants showed an increase in skewness. For the Gamma PDF family that is positively skewed, a negative shift indicates that the tail shrinks (fewer large MMS fluctuations or equivalently, larger IBI averages impacting the scaling in Equation 1). On the other hand, a positive shift indicates a heavier right-tail, with larger MMS fluctuations contributing to this. Given the computation of standardized MMS, *i.e.*, with the denominator containing the average IBI values between the IBI local minima surrounding the IBI maximum, these positive shifts with an increase in MMS then show lower Heart IBI timing values on average. As explained, lower Heart IBI timing values often arise from faster heart rates, which are evident in anxious, stressful states. In this sense, drawing and pointing increase the skewness values (Figure 5 F and G respectively) while the Peg Task includes some participants that increase and others that decrease the skewness during the pain block, relative to the control block (Figure 5H).

The Gamma NSR also shifts in each task relative to rest. In the control block, the drawing task shows a balanced shift as some participants decrease and others increase the NSR, relative to the resting task for both the control and the pain conditions. During the Pointing Task the control case has a spread higher than during pain, and the scatter is biased towards the negative shift. In this case, negative means that the NSR decreased during the drawing task in both the control and the pain blocks, relative to the resting task. Lastly, the Peg Task shows a negative shift during the control block but a positive shift during the pain block. This indicates that relative to the Resting Task, for most participants, the Peg Task evokes higher variability in the IBI fluctuations. Given the relationship in Figure 5A-D between the Gamma NSR and the Gamma shape, and given that the higher the noise, the more randomness in the process, it is safe to infer that as in the Drawing Task, during the pain state, most participants in the Peg Task have more erratic heart rate patterns than during the control state. Interestingly, unlike pointing, which is mediated by kinesthetic (speed and proprioceptive posture feedback) the Drawing Task and the Peg Task are more cognitively demanding. In addition to the speed and posture feedback during the transport phases of the hand motions, these two tasks are also mediated by pressure (haptic) feedback. As such, heart IBI is capturing the increasing difficulties of the tasks: biomechanical goals (Pointing Task), biomechanical and haptic goals (Peg Task), and those plus cognitive / memory goals (Drawing Task).

### 3.3 Tight Linear Relation (Scaling Power Law) Describes the Log Gamma Shape and Log Scale Parameters Fit of the Facial Speed MMS (Micro-Fluctuations)

As we focus on the Gamma NSR, the parameter space spanned by the values along each region forms a three-dimensional scatter that we can also examine across participants for each of the control and pain conditions and across all four tasks. Figure 6 shows the scaling power law relations between the log Gamma NSR and the log Gamma Shape. These comparisons highlight an increase in the noise levels across all three regions of the Face during the pain in the Pointing, the Peg, and the Drawing Tasks, relative to the control condition. Not only do we appreciate a shift in noise patterns in the ophthalmic V1 area, but we also see this shift at the maxillary V2 and the mandibular V3 areas where the noise and the randomness of the moment-by-moment micro-fluctuations in speed deviations from the empirical mean (as measured by the standardized MMS) increased significantly, particularly for the peg task. Supplementary Table 2 shows the linear polynomial fits and goodness of fit parameter values. These were obtained with the Curve Fitting Toolbox of MATLAB R2023b.

**Figure 6.**
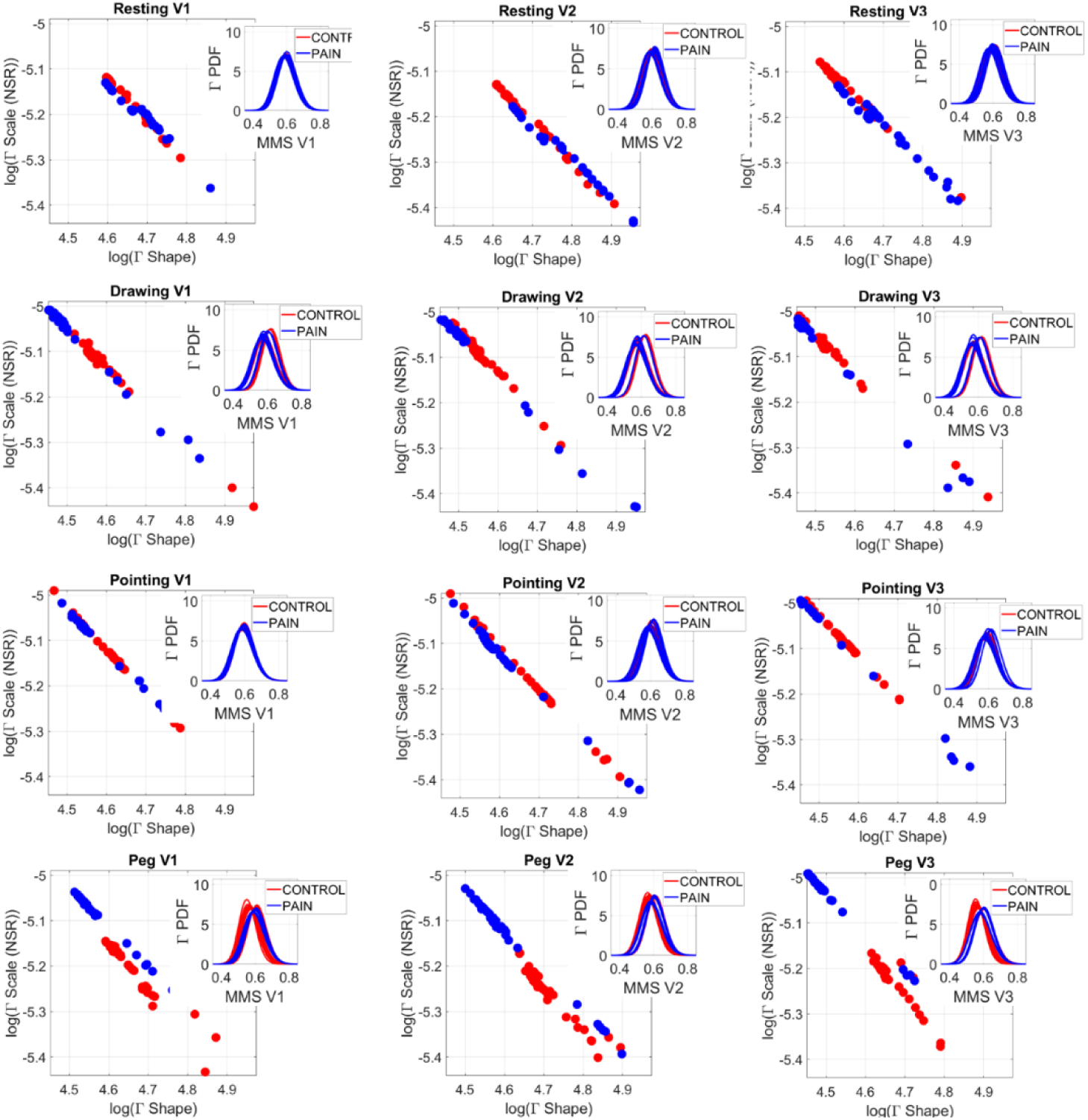
Gamma parameters for facial micro-movements across all regions V1, V2, V3 for all 4 tasks (resting, drawing, pointing and peg) and for control (red dots) and pain (blue dots). Each dot represents a participant empirically estimated (Г_shape_, Г_scale_) measurement, spanning for each participant a family of PDFs. Insets show the Gamma PDFs from the empirically estimated shape and scale parameters. Scatters are well fit by exponential curves and as in the heart IBI case of figure 5, the log-log representation is well fit by a tight linear negative correlation whereby as the Gamma NSR decreases, the Gamma shape increases (see Table 2).

### 3.4 Facial V1 Micro-Movements Shift Across Tasks During Control State

The V1 (ophthalmic) region of the Face comprising the eyebrows, the eyes, and part of the nose, revealed significant differences in the Gamma NSR signatures across tasks, according to the non-parametric Kruskal-Wallis statistical test. This is shown for the control (pre-pain) state in Figure 7A across tasks. Figure 7B shows the within comparisons for the pain condition. The tables in the figure (output by MATLAB) summarize the results in Figure 7C, where we also show the output of the Friedman’s test comparison between the control and pain conditions. Figure 7D shows the pairwise *p-values* obtained from the rank-sum test on the Gamma NSR parameters (equivalently the negatively correlated Gamma shape parameter) across all tasks and conditions for region V1. Figures 6E-F show the pairwise comparisons of the empirically estimated Gamma NSR parameters for V2 and V3, respectively. As can be seen, V1 spanned significantly different families of Gamma PDFs across tasks, while V2 and V3 subregions only show such differences in the tasks relative to the resting task. Given the marked shifts in Gamma PDFs for V1, we next focus on this facial subregion to illustrate the comparisons and examine its relationship with the Heart IBI streams.

**Figure 7.**
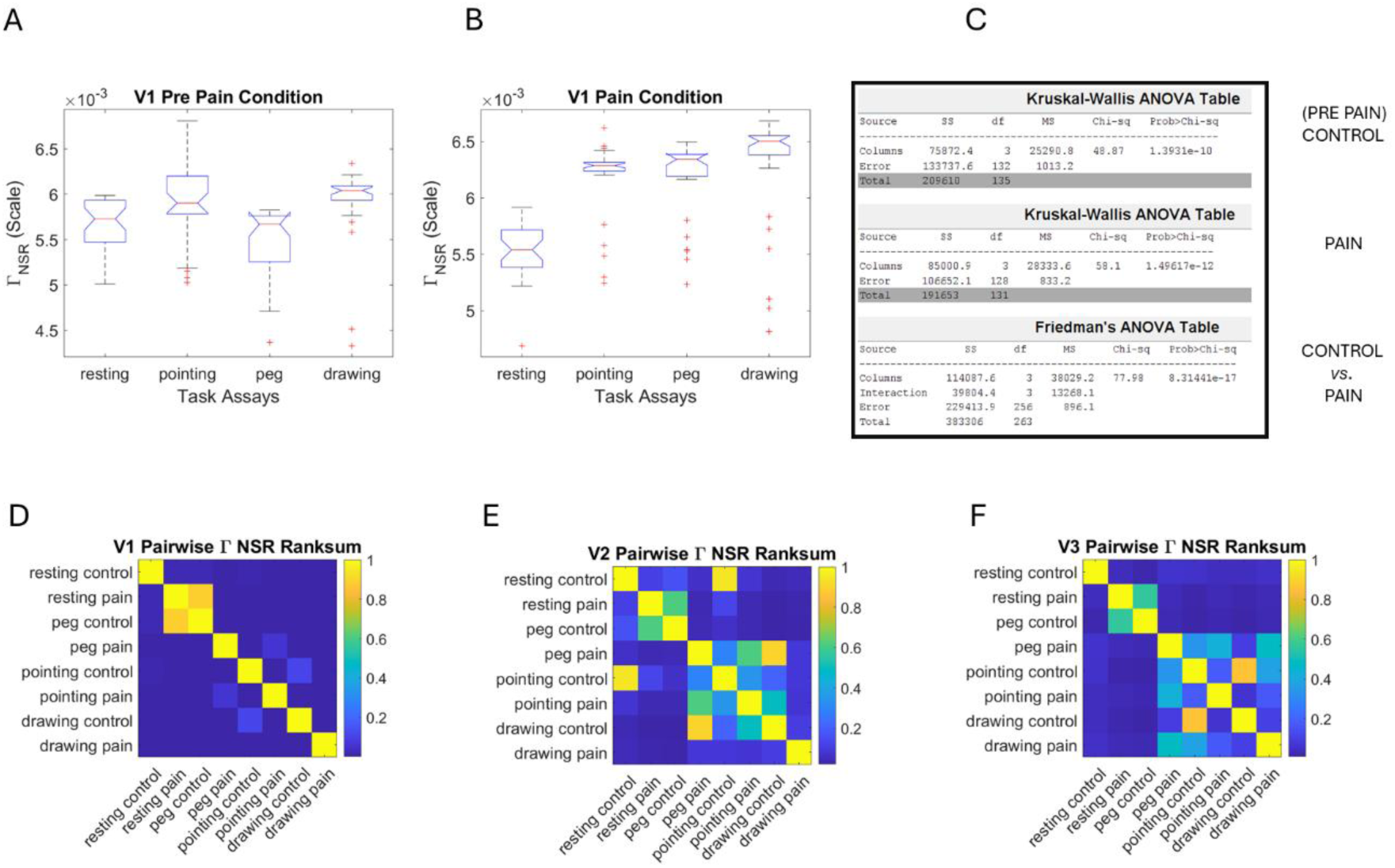
Statistical tests of the Gamma parameters for the facial micro-movements. (A) Control condition (pre-pain) shows statistically significant differences across Gamma NSR parameter according to the non-parametric Kruskal-Wallis test for ophthalmic area V1, across all tasks. MATLAB Multcompare test reveals *p < 0.01* for all tasks pairwise, relative to the resting task. (B) During the pain condition, these differences accentuate. (C) Stats table for comparisons A and B and for the control *vs*. pain comparison, according to the Friedman’s test. All comparisons *p < 10^-10^*. (D) Pairwise comparisons of NSR rank-sum test in ophthalmic area V1, (E) maxillary area V2 and (F) mandibular area V3.

### 3.5 V1 Micro-Movements Shift Across Tasks During Pain States

As with the MMS streams derived from the Heart IBI timings, here we derived the MMS from the V1, V2, and V3 regions of the Face and found the best fitting for the MMS micro-peaks using the continuous Gamma family of PDFs, in an MLE sense. We empirically estimated the shape and the scale parameters with 95% confidence. As we examined the shape and scale relations in Figure 6, we appreciate a shift in the Gamma NSR from the control to the pain states. This shift is most pronounced for the Peg Task and the Drawing Task but also present in the biomechanical Pointing Task. The insets across the panels show the shift as well in the Gamma PDFs derived from the empirically estimated parameters. Notable is a shift upward with the pain condition, within the Pointing, Peg, and Drawing Tasks, across all facial subregions. This increase in noise is also accompanied by a decrease in the shape value, tending towards the memoryless exponential range of the Gamma family (towards the extreme value of 1 denoting randomness). This is like the patterns observed in the IBI signal.

Figure 8A shows the shifts of the Gamma NSR across the V1, V2, and V3 regions represented as points in the 3D parameter space spanned by these triads of values. The scatter shows the increases in noise across the Face, while Figure 8B summarizes these shifts within the parameter plane spanned by the Euclidean norm of the (V1, V2, V3) points taken as vectors from the (0, 0, 0) origin. These are expressed as the scalar magnitude of such vectors for both, the NSR and skewness components across the Face. There we see during the Resting Task, two groups of participants that split values within the pain case, one increasing skewness relative to the control case and the other decreasing it or maintaining it at the lower end of the range of the control values.

**Figure 8.**
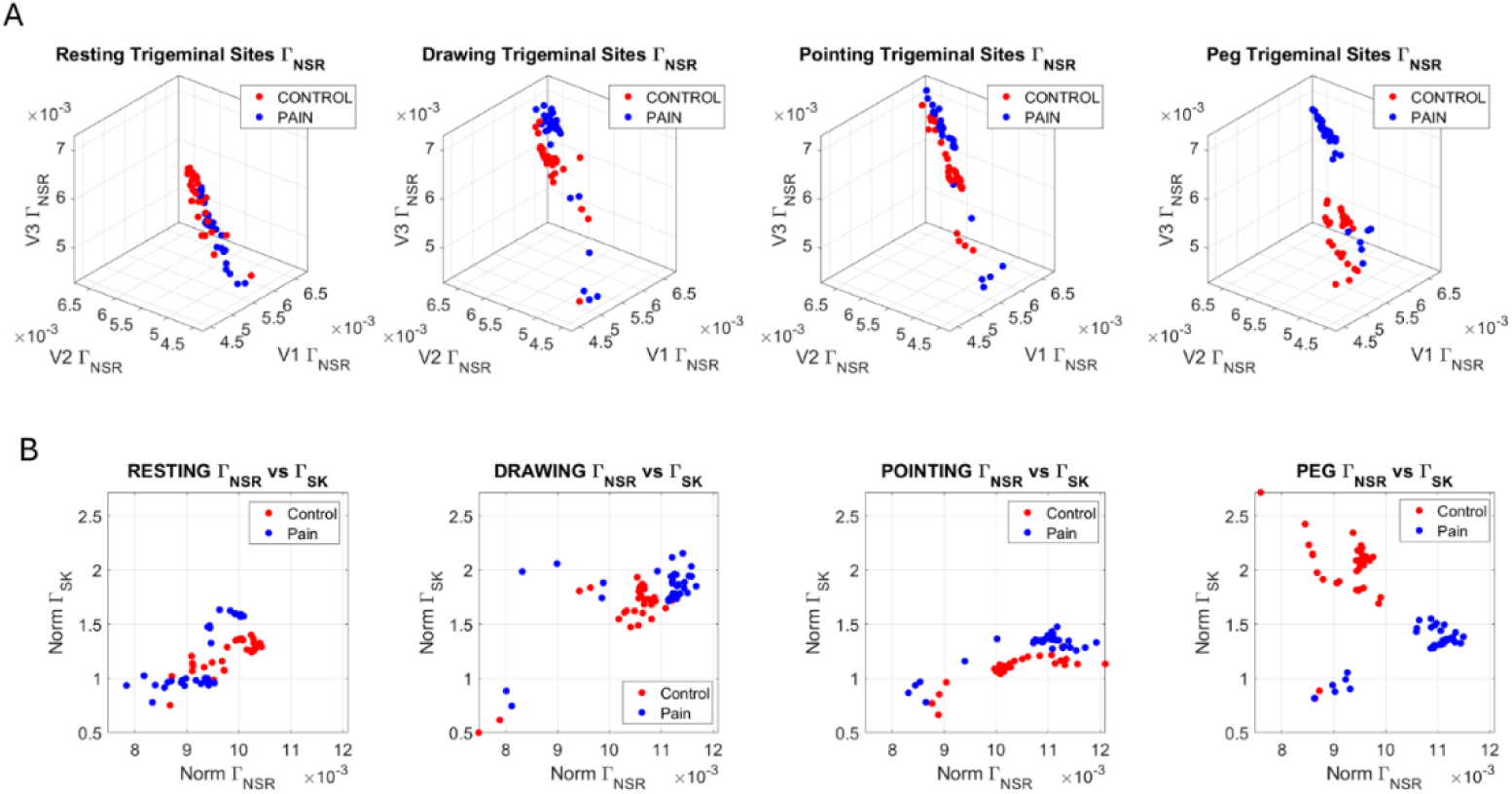
Different representations of the Face data capture the shifts in Gamma families of probability distributions from control to pain conditions and across the different tasks. (A) Parameter space spanned by V1, V2, V3 Gamma NSR with control (red dots) and pain (blue dots) whereby each dot represents a participant, and each parameter space denotes the state of the Face during the task. (B) The Euclidean norm of the parameters vector (V1, V2, V3) is obtained for both the NSR and the skewness and the scalar quantities represented on a parameter plane spanned by the Norm Г_NSR_ and the Г_Skewness_. Notice the different shifting patterns (see details in the results section).

An increase in the skewness indicates a heavier tail of the PDF, suggesting larger values of the MMS. Recalling here Equation (1), the MMS are absolute deviations from the empirically estimated Gamma mean speed amplitude. Larger values of such normalized deviations from the empirical mean occur when the average speed (in the denominator term of the normalizing expression) is smaller, *i.e.*, slower speed. In contrast, lower skewness towards a more symmetric density accumulating lower MMS values occur when the average speed is higher. While pointing in pain, the skewness increases (lower speed of the facial MMS micro-fluctuations on average). In contrast, the skewness decreases during the Peg Task in pain (denoting higher speed of the Face MMS micro-fluctuations on average) and the noise increases (which also brings more randomness with a decrease in shape values). The Drawing Task in pain also increases the noise and randomness levels, while the skewness remains comparable to that of the control condition for most of the participants. These results demonstrate very different patterns for the Peg Task and the Drawing Task involving haptic feedback and a higher cognitive and memory load than for the pointing to a visual target and receiving feedback from the hand speed and the rotational arm joints, as the transport phase of the motion unfolds. In all 3 tasks, pain levels increased relative to rest, but the speed of the facial micro-movements behaved differently.

### 3.6 Distribution Parameters of V1 Region Distinguish Tasks and Conditions

The statistical significance of the pairwise comparisons across the tasks and conditions for the ophthalmic region V1 discussed in Figure 7A-D motivated us to further compare the PDF families in this region. To that end, we used the earth mover’s distance (EMD) (26, 27) and compared pairwise, for each participant, the PDFs generated by the MMS derived from the speed peaks. The results are shown in Figure 9A distinguishing the pre-pain to the during-pain (denoted post-pain) activities for each task. This information was fed to a clustering tree, and 8 subgroups were identified in Figure 9B. Their composition from left to right was 96% control, 57.1% control, 52.3% control, 100% control, 50% pain, 64.2% pain, 100% control and 100% control.

**Figure 9.**
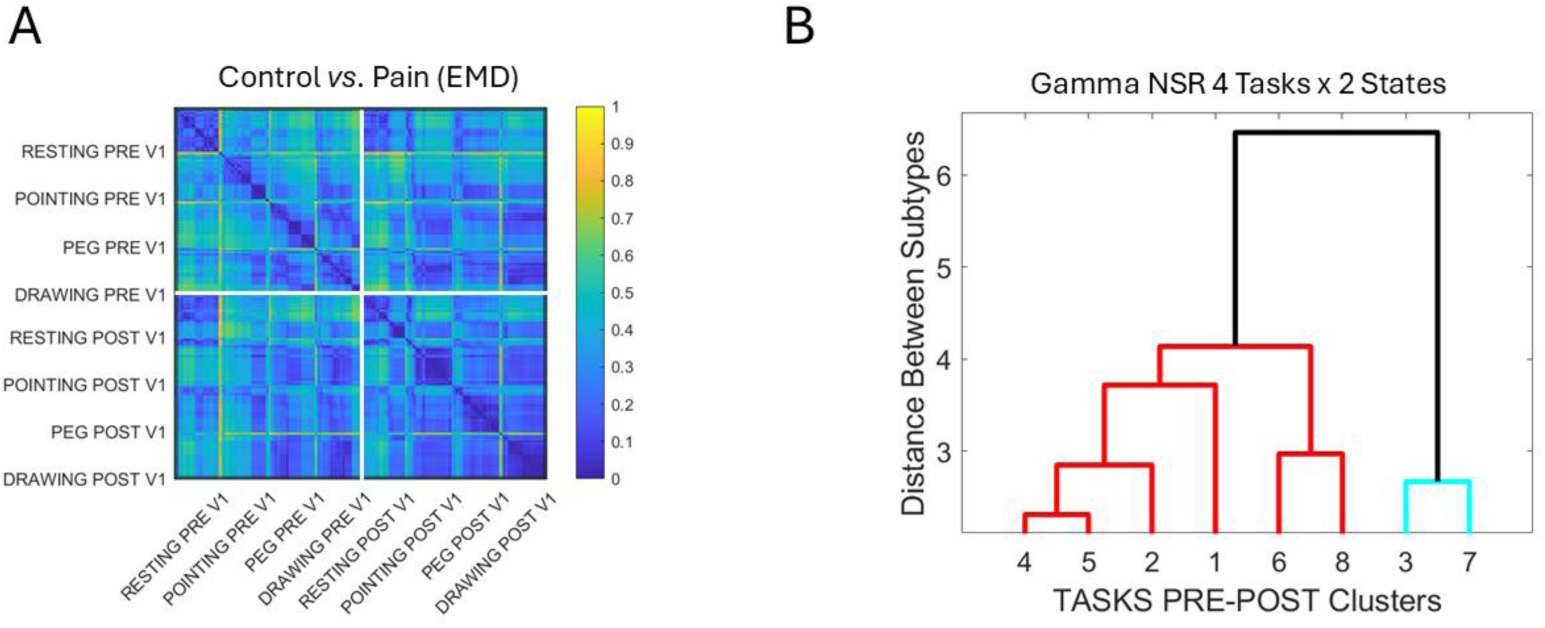
Individualized probability distribution families across participants are compared pairwise using the EMD and grouped into self-emerging clusters. (A) Matrix of EMD values denoting the normalized distance (color bar) for each participant in the V1 area (the most significantly impacted by tasks and pain) – PRE refers to the control (before pain condition) and POST refers to pain the condition. (B) Tree clustering of the participants into 8 groups according to the EMD values in (A) obtained from the distributions of Gamma NSR parameters empirically estimated. Cluster composition of control *vs*. pain PDF percentages are detailed in the main text.

### 3.7 Linear Trend Between NSR Shifts of Face and Heart IBI Control *vs*. Pain

The differences between pain and control Gamma NSR were obtained from the Facial and the Heart’s IBI MMS. Linear polynomial fits were obtained for the scatters of each task, and these revealed a linear trend for the Pointing Task, the Peg Task, and the Drawing Task. These are reported in Supplementary Table 3 and shown in Figure 10.

**Figure 10.**
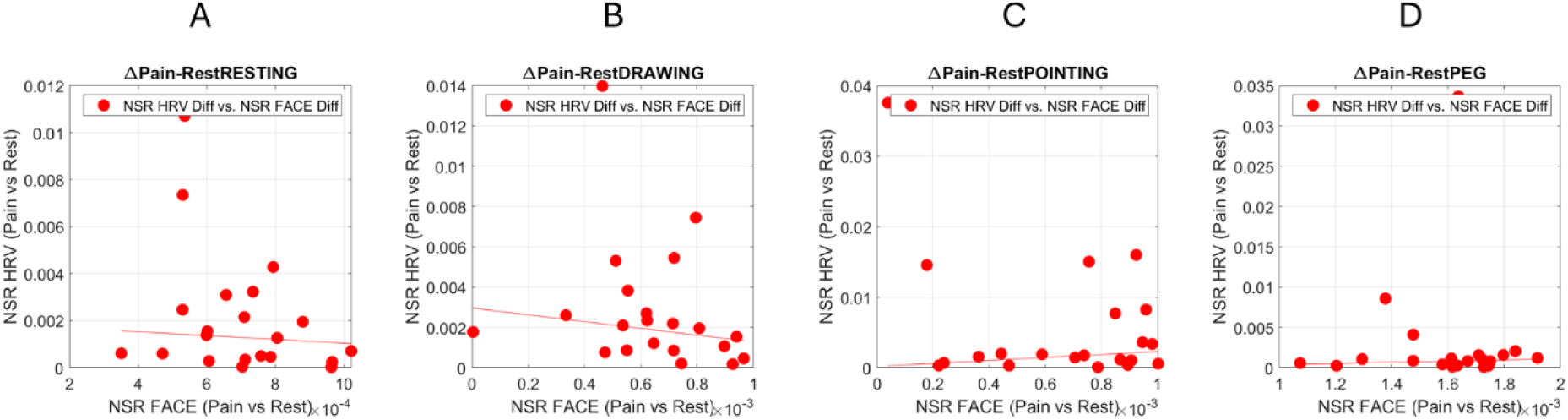
Linear correlations between the difference in Gamma NSR shifting from control resting to the pain condition for each task. (A) Resting task. (B) Drawing task. (C) Pointing task. (D) Peg task. Each point represents a participant. Supplementary Table 3 reports the correlation values and the goodness of fit.

### 3.8 Transfer Entropy: The Face Reflects the Increasingly Erratic Heart Signal with Pain

The TE values for were sorted in increasing order across participants, *i.e.*, for the case when knowing the past IBI & Face signal reduced the Face signal uncertainty, thus making the Face signal more predictable. For each participant then, the TE in the opposite direction, *i.e.*, when knowing the past IBI & Face activity reduces the uncertainty of the Heart IBI activity, was plotted such that we could see for each participant, the deviation of this TE relative to the sorted (from low to high) TE curve.

The resulting curve for the control case (no pain) revealed that in the peg task, during this control condition, there seemed to be an individualized threshold such that those participants with a lower value of TE in the case of *T_IBI_*_→*Face*_ (past heart IBI & FACE activity helps predict Face, red curve) tended to have more pronounced effects of reducing the uncertainty of predicting the heart IBI signal when knowing same past activity, blue curve, than those participants with higher values of *T_IBI_*_→*Face*_. Figure 11A shows the traces for all tasks and points out the Peg Task case. Figure 11B shows that this trend is not present when we invert the comparison.

**Figure 11.**
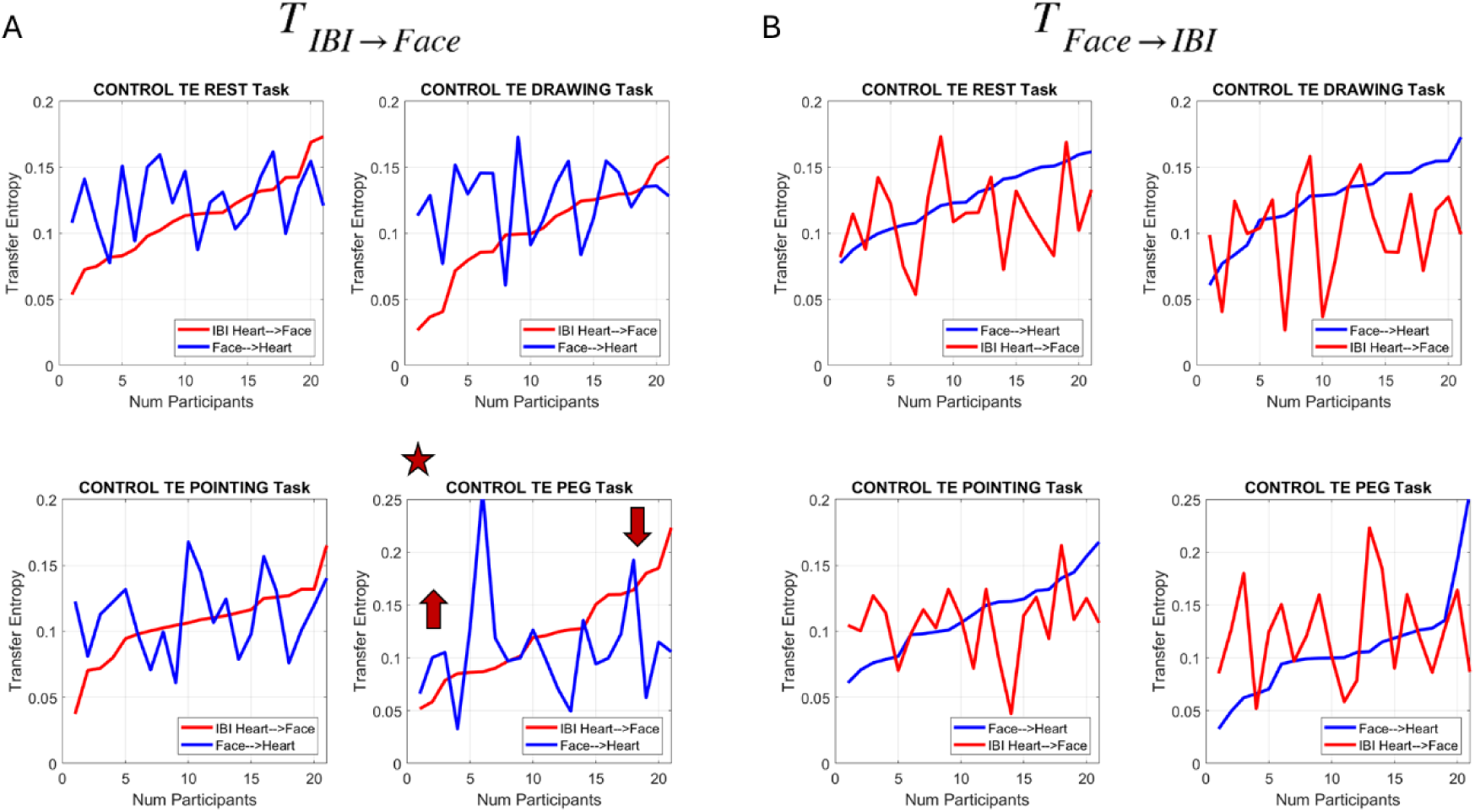
Measures of Transfer Entropy during the Control session for all 4 tasks and across participants. (A) TE sorted in increasing levels within the sample of participants (used as an individualized threshold) for the case where the past heart IBI & FACE activity predicts the current Face activity. Superimposed for each participant is the TE curve (denoted Face → IBI) where past heart IBI & FACE activity predict current heart-IBI activity. The plot with the star highlights the Peg Task where participants with lower values of TE (in IBI → Face) express higher TE (in Face → IBI) for lower thresholds and lower TE (in Face → IBI) for higher thresholds (of IBI → Face TE). (B) Same analysis as in (A) but inverting the process by sorting the Face → IBI TE signal across participants.

Interestingly, the Peg Task was the one with the highest linear correlation (adjusted R-square 0.84 in Supplementary Table 3 and Figure 10) between the Face and the Heart signals capturing the shifts in Gamma NSR from control to pain. This motivated us to perform this analysis for all the tasks during the pain condition. We asked to what extent the pattern observed in the Peg Task would also be observed in the other tasks when the person experienced pressure pain.

Figure 12A reveals that during the pain condition, the effects observed in the control condition for the peg task extend across all 4 tasks. There seems to be an individualized threshold whereby individuals with lower TE (more uncertainty), in the case where past activity of the heart IBI & FACE is predictive of present Face activity, tend to have a higher TE (lower the uncertainty of prediction) for the same past activity predicting present heart IBI activity. Inverting this analysis also shows marked changes relative to the control condition (Figure 12B, where we highlight the peg condition for comparison purposes).

**Figure 12.**
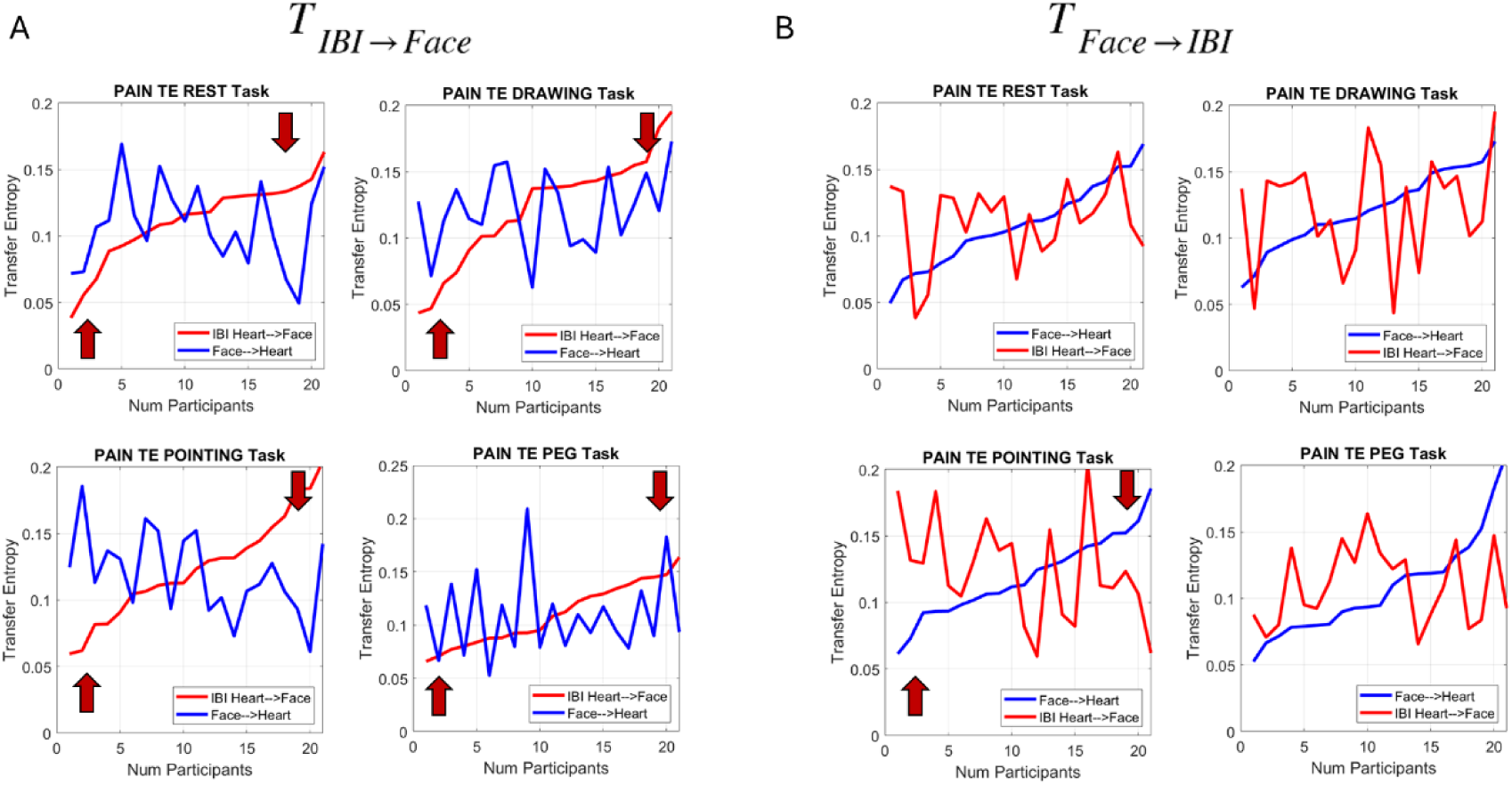
Measures of Transfer Entropy during the Pain session for all 4 tasks and across participants. (A) Identified pattern in the Peg Task shows consistently during the pain state in other tasks. (B) Results from the reverse analysis also show marked differences from the control case (peg task highlighted for comparison purposes).

Given the patterns during the pain condition, we proceeded to obtain the relative differences between the sorted values (smoothly increasing curve) and the varying one, during both the control and the pain cases. The results are presented in Figure 13 in the order in which the experiments took place. We appreciate the overall decrease in TE relative differences during the pain condition (dotted lines) both when the past Face & Heart IBI signals predict the present Heart signal, and when the past Face & Heart IBI signals predict the present Face signal. In both comparisons, the Resting Task and the Drawing Task, we see a higher difference in TE when the uncertainty to predict the present Heart IBI signal decreases, whereas the difference is not as pronounced for the Pointing Task and the Peg Task. Perhaps some possible adaptation or fatigue effect is responsible for this pattern.

**Figure 13.**
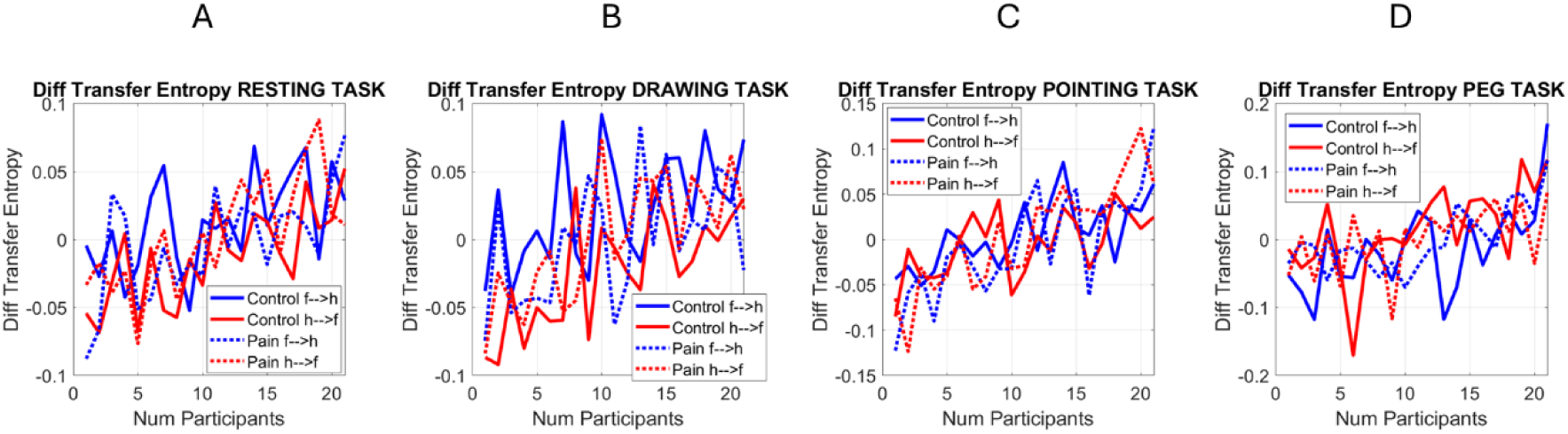
Differences between TE for each of the control and pain conditions, across participants (f→h for Face to Heart and h→f for Heart to Face). (A) Resting Task. (B) Drawing Task. (C) Pointing Task. (D) Peg Task.

Regardless of potential effects of fatigue or adaptation to pain, the Face and Heart activities seem not only correlated but also take turns leading and lagging each other in ways that knowing one in the recent past contributes to reducing the uncertainty of predicting the other in the present. The Heart signals of individuals with more uncertainty in the past Heart IBI (more erratic heart timing) increasingly allows Face signals to show this with higher certainty, a trend that fades away when the uncertainty of the past Heart signal in predicting the Face decreases.

A more regulated Heart in the recent past does not contribute as much as a dysregulated Heart in the prediction of present Face signals. During the pain state, when the Heart signal is more dysregulated, the recent activity of the Heart may be reflected in the present facial micro-movements, suggesting the Face as a proxy of recent Heart dysregulation. We sampled video of the Face with a regular webcam at 30Hz, ∼every 33.3ms, such that in our analysis, 5 frames represent approximately 166.7ms of past Heart activity considered to obtain the TE measure indicating a reduction in the uncertainty of the Face activity. We discuss next the implications of the present results, potential caveats of our study, and next steps in the research.

## 4 Discussion

This work examined the relationship between the Heart IBI signals and the Face micro-motions derived from 10-second-long videos and concurrently recorded in a subset of participants while they engaged in four tasks, during a control condition (pre-pain) and during a pain condition (post experiencing pain). We leveraged recent advances in methods of Artificial Intelligence and Machine Learning to acquire facial kinematics data using the Open Face software (28). These methods provide a grid of points describing the changes in grid pixel point positions over time. By extracting the rate of change of position per unit time, we obtained the speed temporal profiles of 68 points across the grid and studied separately the ophthalmic V1, maxillary V2, and mandibular V3 subregions of the Face in correspondence with the biorhythmic data of the Heart IBI timings.

An empirical characterization of the standardized data type created in our lab and coined the Micro- movements Spikes (MMS, or micro-peaks) was used to represent the moment-to-moment micro-fluctuations of both the Heart IBI peaks and the Face speed peaks, relative to their corresponding (empirically estimated) Gamma mean. In both time series, the biorhythmic micro-fluctuations that they output from the peripheral and autonomic nervous systems were well characterized by the (empirically estimated) continuous family of Gamma probability distributions. The Gamma shape and Gamma scale parameters provided, in both cases, a good representation in an MLE sense, of the peaks of the standardized MMS data. Increases in the Gamma NSR (scale parameter) corresponded to more randomness in the micro-fluctuations, *i.e*., a decrease in the shape parameter, towards the memoryless exponential regime of the Gamma family, where the shape is 1. This relationship was well fit by tight linear polynomial equations on the log-log shape-scale Gamma parameter plane (reported in Supplementary Table 1 for the Heart IBI MMS and in Supplementary Table 2 for the Face MMS data). And since under this tight fit, knowing the Gamma NSR accurately infers the Gamma shape, we focused our analysis on a single parameter, the Gamma NSR, which is the Gamma scale parameter indicating the spread of the distributions. Each participant spanned a family of Gamma PDFs across the tasks (resting, drawing, pointing, and the grooved peg) and conditions (pre-pain or control and pain). We then used this Gamma NSR parameter in combination with another parameter, the Gamma skewness, to lend interpretation to our data.

Furthermore, given this power law description of the parameters of the distribution families that each participant spanned across tasks and conditions, we asked if the two Gamma NSR parameters from the micro-fluctuations of the Heart and the Face streams were correlated. First, we explored the activities of the three regions of the Face and found that the ophthalmic V1 region maximally expressed the shifts in noise signatures from control to pain, across the tasks. We focused on this V1 region then and found statistical significance of these effects.

The shifts in Gamma NSR of the Face V1 region during the pain condition, relative to the control state, were then tested against the shifts in Gamma NSR of the Heart IBI. We found linear correlations with R-squares of 0.84 and 0.85 in the Pointing and the Peg task, respectively (the last two tasks that the participants performed). One aspect of the pain assay may be adaptation, while another aspect of the tasks performed might be fatigue. The fact that the statistical effects were strong at the end of the peg block and the correlation between Heart and Face were stronger in the Peg Task than during the Resting Task or during the earlier Drawing Task, suggests that pain adaptation or fatigue alone could not account for the pain effects quantified here.

To further explore the nature of the relationship between the Heart and the Face signals, we calculated the Transfer Entropy between the two processes, in both directions. This analysis identified an interesting pattern in the Peg Task during the control condition. By sorting the TE values (past activity of the Heart & Face decreasing the uncertainty of predicting the present Face activity), we could see that the TE behaved in an interesting fashion. Those participants with increasingly erratic Heart activity (higher noise and randomness in the micro-peaks fluctuations around the empirically estimated Gamma mean) expressed higher TE for predicting present facial ophthalmic activity. This was specific to the Peg Task in the pre-pain, control condition. How would this finding reflect in other tasks with the Pain condition?

Using the same analysis during the pain condition extended the results to all other tasks. This result suggested a sort of individualized threshold of TE indicating under what conditions the Face V1 activity is predictive of the Heart IBI activity. It appears to be the case that Face activity in the ophthalmic region can act as a proxy of the prediction of an increasingly erratic Heart activity. Intuitively, as pain escalates and the heart reveals it, the Face can broadcast it in the ophthalmic region around the eyes. Here we could quantify this while using the moment-by-moment fluctuations of the standardized MMS (micro peaks) of both the Heart IBI and the Face.

We performed the comparison in the opposite direction and observed congruent results. This motivated us to obtain the absolute differences between the sorted (monotonically increasing) TE values in one direction and the changing TE curve in the other direction. We found that the control condition expressed higher ranges of differentiation between the direction of comparison, with the signal predicting the Face from the dysregulated heart leading in the resting and drawing task but mostly lagging in the pointing and peg tasks. In contrast, the pain condition showed narrower ranges of differentiation across tasks. This suggests that during pain, independently of the task, and under the constraints of our analysis ∼166.7ms into the past, both signals from the Face and heart contribute to the prediction of one another, in a periodic manner (as per the peaks and valleys captured in our data).

An interesting aspect of the results is that the Drawing Task, which has a higher cognitive and memory load (as it demands keeping a memory of the alphanumeric order for random spatial locations of the digits and letters across the board) and must be completed under time constraints (fast), showed less of an effect than the Peg Task. Both tasks had a haptic component whereby the continuous pressure feedback was important. Yet the Peg Task did not have the type of memory load or cognitive demands that the Drawing Task did. In this sense, it is possible that the cognitive and memory loads acted as a distractor of the pressure pain in the Drawing Task, whereas in the Peg Task the haptic (pressure) feedback might have amplified the pain effect. It will be important to explore this proposition with more participants, as this cohort was modest. Furthermore, it will be important to shuffle the tasks and perform the Drawing Task last, so we rule out fatigue effects possibly impacting the pain effects. It is worth noting that the Drawing Task also showed very pronounced pain effects, even though it was performed earlier than the Peg Task. The effects of pain in both tasks were highly separable. If indeed the cognitive and memory demands act as distractors of pressure pain, it will be important to further explore potential therapeutic effects of tasks that while mediated by pressure (haptic) feedback, also have higher memory and cognitive demands in cases where descriptions of pain are verbalized as pressure-like sensations.

Although our results were robust and the effects on the parameters spanning personalized families of PDFs were statistically significant, we caveat that the cohort was modest. In future studies we plan to highly scale the study and test participants longitudinally. This approach will help us further ascertain the individualized (personalized) shifts in Gamma distribution families over time, as participants transition from the Control to the Pain condition. Under such conditions, we will be able to document the pertinent parameter ranges across the population. These will include participants with chronic pain, transient pain, and dysregulated heart activity of various kinds, not necessarily induced by pain. For example, we have found that individuals on the autism spectrum have dysregulated heart IBI by default (4, 5) – during resting state – and this dysregulated state also has corresponding dysregulated facial micro-movements at rest (29). We plan to explore various pathologies of the nervous system to further characterize the connection between the Heart and the Face biorhythms.

The present analyses produce empirical families of PDFs of the moment-to-moment fluctuations present in natural human biorhythmic activities across different systems. We offer here a unifying framework amenable to not only characterize and capture individualized families of distributions and parameter ranges for each person but also do so across the population. This is important information for generative AI models that currently rely on synthetically produced data from artificially generated distributions, or even mixed data from natural sets. By adopting our methods, it will be possible to generate empirically validated biological variations of time series data registered from the human nervous systems. This approach can counter Model Autophagy Disorder (MAD) (30) where generative AI models degrade performance when trained in synthetic data generated by previous iterations of the same model. There is no need to do so when a single individual generates entire families of distributions capable of differentiating across tasks and different bodily and mental states (*e.g.*, pain *vs.* pain-free states).

In conclusion, the assays employed here are simple and brief, offering new multimodal means to quantify the effects of pain experiences using our new statistical platform for individualized behavioral analysis (SPIBA) (31). This work offers new non-invasive measures such as short Face videos to predict dysregulation of Heart IBI. More erratic IBI seems to be highly predictive of corresponding Face micro-motions, particularly those of the ophthalmic V1 region where using our approach, one might be able to see more than meets the eye.

## 6 Author Contributions

Conceptualization (EBT and ME), Data curation (EBT and ME), Formal analysis (EBT and ME), Funding acquisition (EBT), Investigation (EM), Methodology (EBT and ME), Project administration (EBT), Resources (EBT), Software (EBT), Supervision (EBT), Validation (EBT and ME), Visualization (EBT and ME), Writing original draft (EBT), Writing, review and editing (EBT and ME).

## 8 Acknowledgments

We thank the participants and the members of the Sensory Motor Integration Laboratory.

## 9 Data Availability Statement

The datasets generated for this study are available at https://zenodo.org/records/17082141.

