## Supplementary Tables for "Facial Micro-Movements as a Proxy of Increasingly Erratic Heart Rate Variability While Experiencing Pressure Pain"

### 1 Supplementary Tables

Supplementary Table 1. Polynomial fitting values of the heart IBI MMS according to the empirically estimated Gamma parameters (21 participants). Gamma plane log-log scatters well fit by

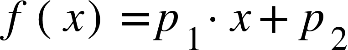
 for slope and intercept values without centering the data. Mean and SD of centered data are also reported.

| Task - Condition | R-square | Adjusted  R-square | SSE | RMSE | Slope *p_1_* | Intercept *p_2_* | Centering |
| --- | --- | --- | --- | --- | --- | --- | --- |
| *Resting Control* | .999 | .999 | .0140 | .0272 | -.9874 | -.505 | Mean 3.23  Std .887 |
| *Resting Pain* | .998 | .998 | .0249 | .0362 | -.9854 | -.500 | Mean 3.50  Std .827 |
| *Drawing Control* | .998 | .998 | .0146 | .0277 | -.9988 | -.448 | Mean 3.38  Std .821 |
| *Drawing Pain* | .998 | .998 | .0145 | .0276 | -.9911 | -.489 | Mean 3.71  Std .641 |
| *Pointing Control* | .998 | .998 | .023 | .0349 | -1.0003 | -.459 | Mean 3.45  Std .827 |
| *Pointing Pain* | .995 | .994 | .025 | .0367 | -.9861 | -.511 | Mean 3.44  Std 0.518 |
| *Peg Control* | .999 | .999 | .010 | .0235 | -.9983 | -.466 | Mean 3.78  Std 0.999 |
| *Peg*  *Pain* | .998 | .998 | .024 | .0355 | -.9812 | -.524 | Mean 3.71  Std 0.647 |

Supplementary Table 2. Polynomial fitting values of the face MMS according to the empirically estimated Gamma parameters (36 participants). Gamma plane log-log scatter well fit by

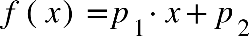
 slopes and intercept values without centering the data. Mean and SD of centered data are also shown.

| Task - Condition | R-square | Adjusted  R-square | SSE | RMSE | Slope *p_1_* | Intercept *p_2_* | Centering |
| --- | --- | --- | --- | --- | --- | --- | --- |
| *Resting Control* | .9941 | .9939 | .00047 | .00383 | -.9303 | -.8410 | Mean 4.655  Std .0530 |
| *Resting Pain* | .9647 | .9636 | .00234 | .00870 | -.7767 | -1.5611 | Mean 4.682  Std .0577 |
| *Drawing Control* | .9938 | .9936 | .00122 | .00601 | -.8264 | -1.3340 | Mean 4.593  Std 0.0912 |
| *Drawing Pain* | .9919 | .9916 | .00192 | .00752 | -.8743 | -1.1140 | Mean 4.522  Std .09394 |
| *Pointing Control* | .9991 | .9916 | .00012 | .00192 | -.9350 | -.8194 | Mean 4.611  Std .0713 |
| *Pointing Pain* | .9976 | .9976 | .00025 | .00271 | -.8743 | -1.0987 | Mean 4.563  Std .0634 |
| *Peg Control* | .9355 | .9336 | .00986 | .01679 | -.9182 | -.9298 | Mean 4.654  Std .0686 |
| *Peg*  *Pain* | .9985 | .9984 | .00015 | .00212 | -.8713 | -1.1059 | Mean 4.565  Std .0626 |

Supplementary Table 3. Linear trends between the shifts in Gamma NSR of Face and IBI from Control to Pain conditions

| Task | R-square | Adjusted  R-square | SSE | RMSE | Slope *p_1_* | Intercept *p_2_* |
| --- | --- | --- | --- | --- | --- | --- |
| *Resting* | .4349 | .4080 | .0015 | .0085 | -.0009 | -.0092 |
| *Drawing* | .7673 | .7562 | .0024 | .0107 | -1.3061 | .0089 |
| *Pointing* | .8425 | .8350 | .0050 | .0155 | .0003 | .0038 |
| *Peg* | .8510 | .8439 | .0030 | .0120 | -1.6474 | .0070 |
